# Overexpression of *Medicago sativa glutamate-semialdehyde aminotransferase* (GSA) gene in tobacco increased photosynthesis efficiency

**DOI:** 10.1101/640425

**Authors:** Maryam Ghasemzadeh, Mahdi Khozeai, Hamzeh Amiri

## Abstract

To investigate the effect of increased *glutamate-semialdehyde aminotransferase (GSA)* on photosynthetic capacity and growth, tobacco (*Nicoliana tabacum* L. Xanti) plants with increased levels of glutamate-semialdehyde aminotransferase protein were produced. This was achieved using a cassette composed of a full-length *Medicago sative* cDNA under the control of the cauliflower mosaic virus 35S promoter. The results revealed distinct impacts of GSA activity on photosynthesis rate and growth in *GSA* over expression tobacco plants. In transgenic plants with increased GSA activity, an increase in soluble and insoluble sugars accumulation was evident. Total biomass, leaf area, plant height and internode 3-4 were increased in *GSA* sense plants, compared with equivalent wild-type tobacco plants. Moreover, transgenic tobacco plants with increased GSA activity exhibit higher levels of 5-aminolevulinic acid (ALA) accumulation and increased in content of chlorophyll and carotenoids pigments. Collectively, our data suggest that higher level of GSA activity gives an advantage to photosynthesis, growth in tobacco plants. This work also provides a case study that an individual enzyme in the biosynthesis of chlorophyll pathway may serve as a useful target for genetic engineering to improve photosynthesis and growth in plants.

**Highlight:** Overexpression of *glutamate-semialdehyde aminotransferase (GSA) increase* photosynthetic capacity, growth in tobacco.

## Introduction

5-Aminolevulinate compound is an important biosynthetic precursor of all tetrapyrrole compounds such as chlorophyll, heme, sirohem, cobalamin (vitamin B12), and phytochrom (Fig. 1)(Mochizuki *et al.*, 2010). ALA has received wide attention as a new plant growth regulator (PGRs) and growth-promoter (Bindu and Vivekanandan, 1998), stimulator assimilation of nitrogen (Wei *et al.*, 2012) and sulfur (Maruyama-Nakashita *et al.*, 2010), and as a biodegradable herbicide or insecticide (Rebeiz *et al.*, 1988a; Sasaki *et al.*, 2002; Yang *et al.*, 2011; Xu *et al.*, 2015) in agricultural application. ALA also has medical applications for cancer photodynamic therapy, tumor diagnosis (Wachowska *et al.*, 2011), antimicrobial drug (Banerjee *et al.*, 2010) and improvement sleep (Perez *et al.*, 2013) and other clinical uses. At low concentrations (30–100 mg 1^-1^), ALA increases the yield of agricultural products also can improve plant tolerance to various stresses (Wang *et al.*, 2004, 2010; Sun *et al.*, 2009; Han *et al.*, 2014; Yang *et al.*, 2014; An *et al.*, 2016b,a; Wu *et al.*, 2018) promoting antioxidant activity (Nishihara *et al.*, 2003) but at high concentrations known as an environmental friendly and biodegradable herbicide or insecticide (Jung *et al.*, 2004).

**Fig. 1.**
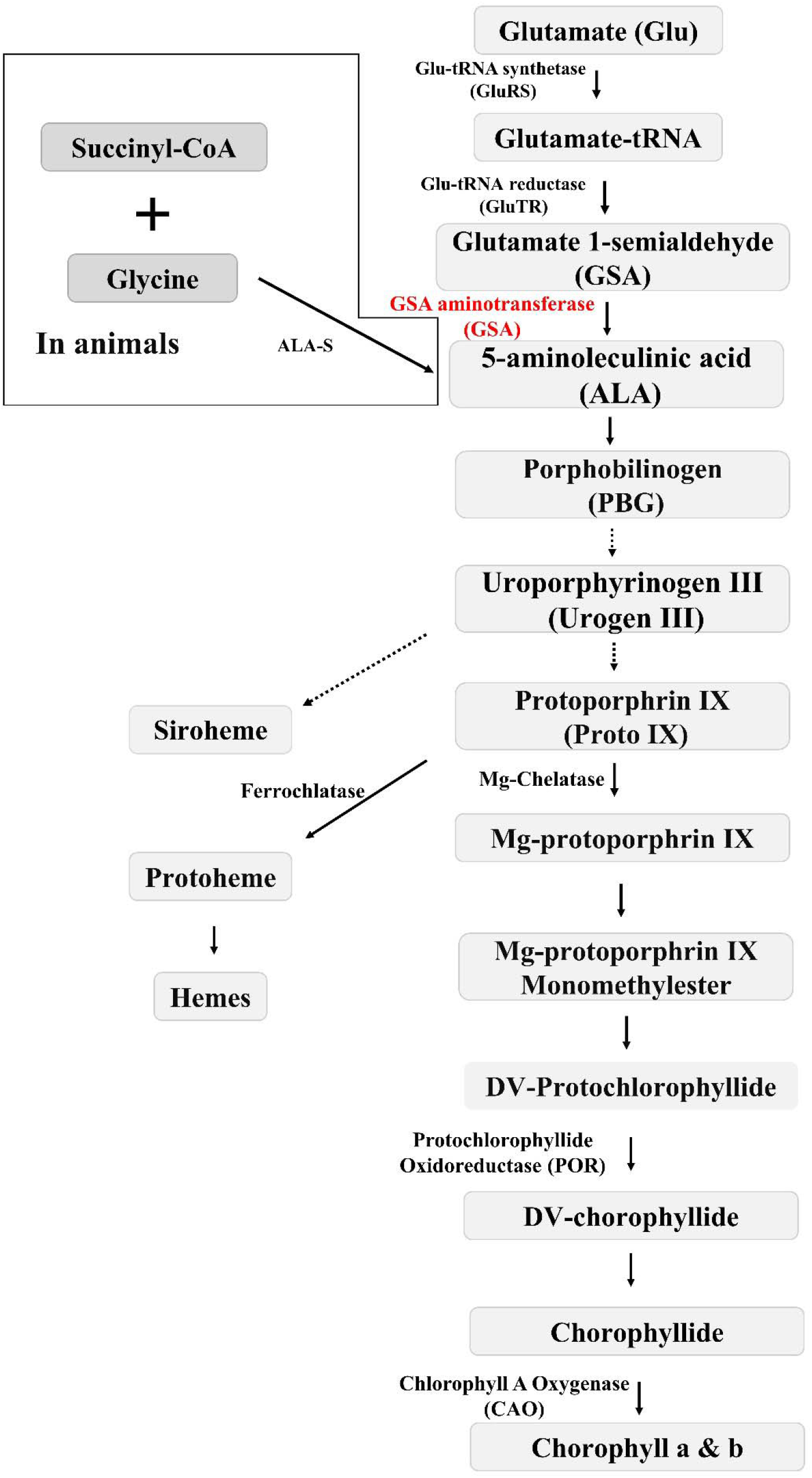
In animals ALA is synthesized from combination of succinyl-CoA and glycine catalyzed by ALA-S, but in Plants ALA is synthesized from Glu that respectively convert by Glu-RS, Glu-TR, and *GSA* to ALA. Two molecules of ALA form PBG then four molecules of PBG are condensed to form Urogen III. Proto IX is synthesis by several modifications in Urogen III. Fe+^2^ and Mg^2+^ insert by Ferroclatase and Mg-Chelatase into Proto IX. The pathway is branched into the synthesis of Heme and phytochrome or Chl a and Chl b at this step. (Dashed arrows indicated several reaction is down and filed arrows indicated one reaction is down).

Two different pathways of ALA biosynthesis in living organisms have been observed. One is the C4 route that known as Shemin pathway. In this pathway ALA is formed from the condensation of succinyl-CoA and glycine, catalyzed by ALA synthetase (ALA-S) in the mitochondria of animals, yeast, fungi or in some kind of bacterial species. Another is the C5 route that known as Beale pathway, in that route glutamate is used as the precursor to form ALA after three enzymatic reactions occurring in the stroma of plastids of higher plants and in bryophytes, cyanobacteria and many eubacteria. Glutamate respectively convert by glutamyl-tRNA synthetase (Glu-RS), glutamyl-tRNA hydrogenase (Glu-TR), and glutamate-1-semialdehyde aminotransferase (GSR) to ALA (Beale, 1990; Wettstein *et al.*, 1995; Iida *et al.*, 2002). Two molecules of ALA are condensed to form the monopyrrole (porphobilinogen; PBG). four molecules of PBG are polymerized to form the cyclic tetrapyrrole uroporphyrinogen III (Urogen III). The pathway is branched at this step to form siroheme (cofactor of nitrite and sulfite reductases which function in nitrogen and sulfur assimilation). Protoporphyrin IX (Proto IX) is formed by further steps including decarboxylations and oxidation. Ferroclatase and chelatase insert Fe^2+^ or Mg^2+^ to Proto IX. Subsequent changes lead to heme or chlorophyll (Chl) or, respectively (Fig. 1).

Chemical production of ALA is very expensive so some studies were down in micro-organisms to biological produce of ALA for medical or agriculture purposes and a few studies were focused on plants (Xie *et al.*, 2003; Choi *et al.*, 2008; Yu *et al.*, 2015; Yang *et al.*, 2016; Kang *et al.*, 2017; Zhang and Ye, 2018).

Up to now, there are many studies shown that exogenous ALA in low concentrations can improve tolerance of plants in stress (Wang *et al.*, 2004, 2010; Sun *et al.*, 2009; Han *et al.*, 2014; Yang *et al.*, 2014; An *et al.*, 2016b,a; Wu *et al.*, 2018; Phour *et al.*, 2018) and as a plant growth regulator (PGRs) are widely applied to improve photosynthesis, growth, development, productivity (Xu *et al.*, 2011; An *et al.*, 2016a,b; Ye *et al.*, 2016; Anjum *et al.*, 2016; Tang *et al.*, 2017; Liu *et al.*, 2018) and good appearance and quality of products (Fuli Xu *et al.*, 2012) of plants under both normal and stressful conditions.

In 1997 the yeast *Hem1* gene (encoding aminolevulinate synthase in C4 route) was transferred into tobacco, The *Hem1* gene could be expressed in transgenic tobacco, so more endogenous ALA was biosynthesized (Zavgorodnyaya *et al.*, 1997). Then, *ALA-S* gene of *Bradyrhizobium japonicum* was introduced into the genome of rice. All transgenic rice lines expressed *ALA-S,* so higher levels of endogenous ALA and tetrapyrrole compounds was achieved (Jung *et al.*, 2004), But transgenic rice lines were grown only under low light condition which was due to the accumulation of protoporphyrin IX, an intermediate of ALA metabolism, in the transgenic rice and photodynamic damage of transgenic lines in high light condition (Jung *et al.*, 2004, 2008). Plant height, tiller number and yield of transgenic rice decreased compared to WT rice (Kuk, 2007). In contrast, overexpression of *Hem1* gene from yeast (McCormac *et al.*, 2001) in to *Arabidopsis thaliana* showed more growth than the WT plants under light condition of PPFD 2 1 about 1000 μmol m^-2^ s^-1^ at noon without any photobleaching (Zhang *et al.*, 2010). The same results also obtained in tobacco transgenic lines with over expression of yeast *Hem1* genes, the transgenic plants with upregulation of *Hem1* gene had higher photosynthetic capacity than wild type tobacco (Zhang *et al.*, 2011). Down regulation of ALA showed growth retardation and leaves phenotype which was accompanied with chlorophyll variegation pattern in antisense transgenic tobacco plants (Höfgen *et al.*, 1994). Genetic manipulation of chlorophyll biosynthesis pathway in Arabidopsis, tobacco, Brassica and other crop plants showed significant changes in the amount of chlorophyll and pigments, and effects on quenching of O_2_ and other reactive oxygen species leads to changes photosynthesis, plant productivity and grain yield in normal growth conditions as well as in a stressful environment (Papenbrock *et al.*, 2000; Tanaka *et al.*, 2001; Alawady and Grimm, 2005).

In this study, *Medicago sativa GSA* was introduced into the genome of tobacco to modulate GSA activity. The results clearly demonstrated that overexpression of *GSA* met our hypothesis regarding to increase photosynthesis rate or positive impact on growth analysis. To the best of our knowledge, this is the first report that genetic engineering of *Medicago sativa GSA* could improve growth and photosynthetic pigments in transgenic plants. This would open up a new way to have more investigation on secondary metabolites and intermediate sugars to study the primary and secondary pathway in transgenic lines.

## MATERIALS AND METHODS

### Preparation Overexpression Plant

In this research, the cDNA of the *GSA* gene in *Medicago sative* L. (accession No: HQ244440.1) (Ferradini *et al.*, 2011) were amplified by PCR using RB pfu polymerase kite (RNA biotechnology, Co, Isfahan Iran) and the following primers; Forward: <u>GGATCC</u>AAAATGGCTGCTTCGGGTATT, Backward: <u>GAGCTC</u>TCAGATCTCCCTAAAGA. The PCR reactions were as follow: for 94 °C for 5 min hot start, 30 cycles of 94 °C for 30 sec denature, 55 °C for 30 sec annealing, 72 °C for 1.5 min extension, and 72 °C for 10 min final extension. pBI121 plant expression vector as a binary vector containing the cauliflower mosaic virus promoter CaMV 35S and transcription terminator sequences (NOS) and ß-glucuronidase gene and a kanamycin resistance gene (NPTII) as a selection marker were used. After the sequence integrity of amplified *GSA* was verified by sequencing, pBI121 and the fragments of *GSA* was digested with Bam HI and Sac I, then the fragments of *GSA* were ligated into pBI121 by T4 DNA ligase. The transfer of the pBI121-GSA was confirmed by PCR colony, digestion and DNA sequencing. Then pBI121-GSA was introduced into *Agrobacterium tumefaciens* strain LBA4404 based on a freeze/thaw protocol (Green and Sambrook, 2012). Then this transformed Agrobacterium used for transformation of tobacco based on Leaf-disk protocol (Horsch *et al.*, 1985). The seventeen primary transgenic Tobacco lines (T0 generation) were selected on kanamycin-containing MS medium (Murasnige and Skoog, 1962) then were rooted on kanamycin-containing medium, then transferred to soil and grown to maturity. In order to confirm the presence of the *GSA* gene in putative transgenic tobacco, analysis of total DNA-PCR, RT-PCR and real time PCR were performed. Based on these selections in the T0 generation, three GSA-overexpressing lines (T0-1, T0-3, and T0-4) were chosen for subsequent analysis. Analysis done in 8-10 leave stage of T0 generation tobacco. Leaf of this lines rapidly frozen in liquid nitrogen for analysis.

T0-1, T0-3, T0-4, and WT Tobacco plants were grown in a controlled-environment cabinet, were Irrigated by Hoagland (Hoagland and Arnon, 1950) (temperature 25±5 °C, relative humidity 60±10 %, PPFD of about 300-350 μmol m^-2^ s^-1^).

### Real-time Quantitative PCR

Real-time PCR was carried out using RB SYBR master mix (RNA Biotechnology, Isfahan, Iran). The total RNA was isolated from fresh leaves using Iraizol reagent (RNA biotech, Iran). cDNA was synthesized using the RB M-MLV reverse transcriptase (RNA biotechnology, Co, Isfahan, Iran). The sequences of the real time primers designed to amplify *GSA* were: GATTCCGTCAAAGGTGCTCG as forward and GTGGCAGCTTTAGGAACACC as Backward. The PCR conditions were 94 °C for 4 min followed by 40 cycles of 94 °C for 10 s, 62 °C for 40 s, 72 °C for 60 s, followed by 7 min at 72 °C. Serial dilutions of cDNA were used to obtain optimized standard curve amplification efficiency and the best cDNA concentration for real-time PCR. PCR was performed in triplicate for each sample, and the expression levels were normalized to that of a tobacco actine gene.

### Morphologic information survey

To calculate total leaf area fifth copper leave for each plant were used in triplicate. Sum of second, third and fourth upper leave used as total dry weight. For measurements of water content, the fourth leaves were weighed. Next, the leaves were dried in an oven at 60 °C for 24h and were weighed. WC was calculated as shown in this Equation.

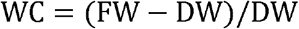

### Photosynthetic pigments content

Arnon (1967) method was used to measure chlorophyll contents (Arnon, 1967). 100 mg of tobacco leaves were homogenized in 2 ml 80% acetone. Extracts were centrifuged at 3,000 g and the absorbance of supernatant was measured in triplicate. Absorbance was measured at wavelengths of 480,645 and 663 nm by ELISA reader (BioTek spectrophotometre Epoch). Chlorophyll content was calculated using the formula of Arnon and expressed in mg g^-1^ fresh weight (FW). Carotenoid content was estimated using the formula of Kirk and Allen (1965) and expressed in mg g^-1^ fresh weight (V: volume of supernatant (ml), W: sample mass (g)) (Kirk and Allen, 1965).

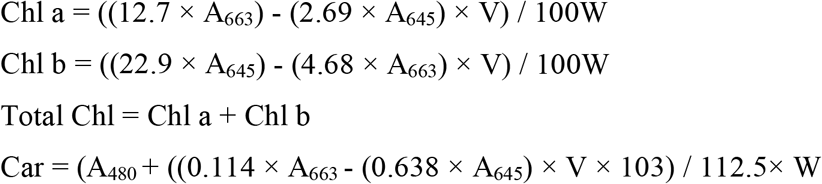

### ALA content

was measured in tobacco plants as described by Mauzeral and Granick (1956) with some modification (Mauzerall and Cranick, 1956) by Khozaei et al (2010). 100 mg of tobacco leaves were homogenized, in 1 ml of Phosphate buffer 50 Mm made by mixing K_2_HPO_4_ and KH_2_PO_4_, pH:6.8. Samples were centrifuged for 10 min at 16000 g, and then 400 μl of supernatant was mixed with 100 μl of ethyl acetoacetate and boiled for 10 min at 100 °C. After cooling the samples were spun again for 5 min at 16000 g and the supernatant was transferred to a new tube and mixed with 500 μl of modified Ehrlich’s reagent (373 ml acetic acid, 90 ml 70% (v/v)perchloric acid, 1.5 gr HgCl_2_, 9.1 gr 4-dimethlaminobenzaldehyde, built reagent to 1 lit adding 500 ml H_2_O). Calorimetric Determination was measured at 553 nm and the ALA content of tobacco samples were calculated using a standard curve generated by commercial ALA. Absorption was measured in triplicate.

### Soluble and Insoluble Sugar Content

Were extracted using the method described by Sheligl (1986). About 100 mg of dried tobacco leaves were extracted in 15 ml of hot 80 % ethanol (80 °C). 1 ml 5 % (w/v) phenol and 5ml concentrated H_2_SO_4_ were added to 2 ml plant extract and mixed thoroughly. The reaction mixture was allowed to stand for 30 min before the absorbance was recorded at 485. Soluble sugars content of the sample was calculated based on calibration curve from a glucose standard. Starch content was extracted from the residual plant material from the soluble sugar extraction described above. Barium hydroxide and Zinc sulfate solvents used for separation of pigments and other components from the residual plant material (Sheligl, 1986). The insoluble products were assayed by the same phenol-sulfuric method described above.

### Photosynthesis Parameters Survey

Gas exchange characteristics [Net photosynthesis (A_n_), and intercellular CO_2_ concentration (Ci)] were measured by portable photosynthesis system (CI-340 Handheld Photosynthesis System-USA) equipped with a 25 mm×25 mm square leaf chamber. Young expanded leaves (second, third and fourth leaf) were measured at light intensity about 300-350 μmol m^-2^ s^-1^ at air temperature about 25 °C and atmosphere CO_2_ concentrations about 380 μl l^-1^.

### Statistical Analysis

SPSS22 software (version 22) was used for statistical analysis. Shapiro–Wilk test used for normality test of data. One-way analysis of variance (ANOVA) and Duncan’s multiple comparison test applied for compare means. Excel was used to draw graphs.

## RESULTS

### Production of Transgenic Tobacco Overexpressing *GSA*

A full-length *GSA* cDNA was used to prepare a sense gene construct in the binary vector pBI121 containing CaMV 35S promoter and NOS terminator (Fig. 2).

**Fig. 2.**
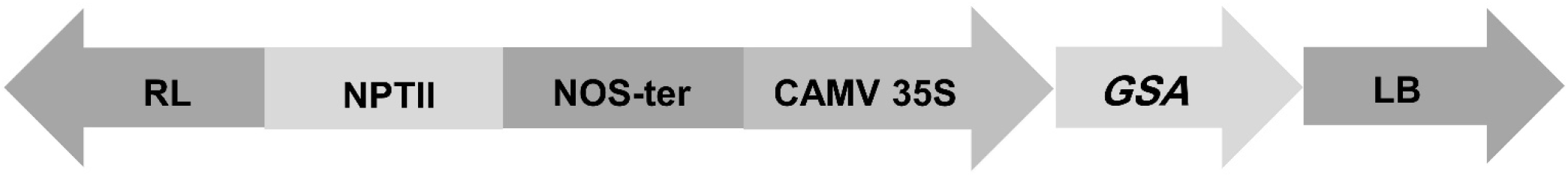
Structure of T-DNA region of binary vector pBI121-GSA constructed for expression of *GSA;* a cassette composed of a full-length *Medicago sative* cDNA under the control of the cauliflower mosaic virus 35S promoter, NOS terminator (NOS-ter) and a kanamycin resistance gene (NPTII) as a selection marker.

The recombinant vector pBI121-GSA was transferred to *Agrobacterium tumefaciens,* and this was used to transform wild-type tobacco plant. Primary transform lines (T0) were selected on kanamycin-containing medium and subsequently moved to soil and grown to maturity, after confirmation of *GSA* gene in the primary resistance lines by PCR and specific primers, expression of the *GSA* mRNA in the kanamycin-resistant plants was confirmed by RT-PCR and concurrently, the quantification expression of *GSA* was carried out by real-time PCR among 17 individual transgenic lines (Fig. 3A).

**Fig. 3.**
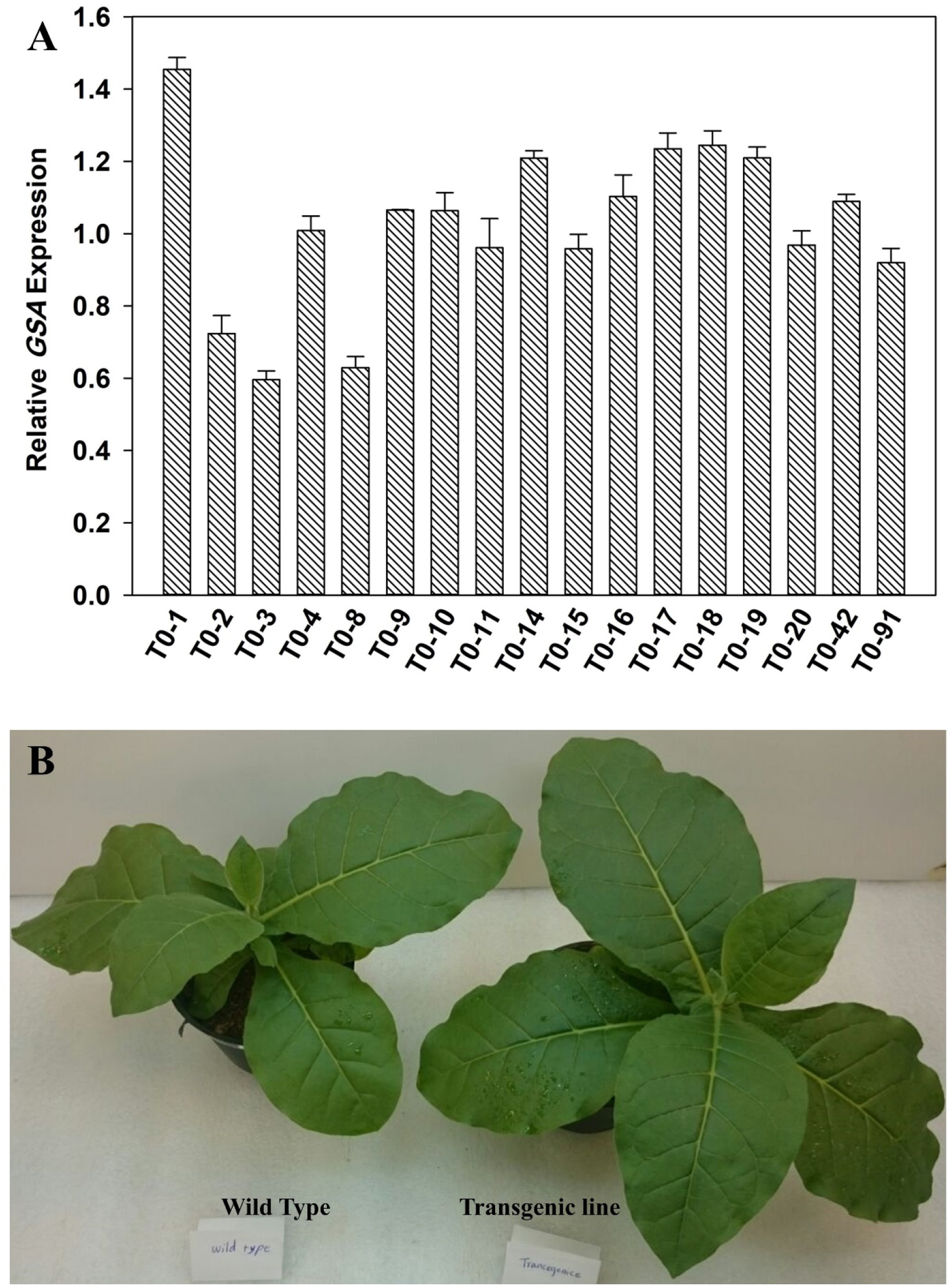
(A) Relative Expression of seventeen line of transforms tobacco. (B) Leaf area of transforms tobacco and wild-type plants are compared. Tobacco plants were grown in a controlled-environment cabinet (temperature 25±5 °C, relative humidity 60±10 %, PPFD of about 300-350 μmol m^-2^ s^-1^).

Based on the gene activity screens in the T0 generation, three *GSA* overexpressing (*GSA*ox) lines (−1, −3, and −4) were selected for further analysis and propagated by self through to the T3 generation.

### Increased GSA Activity Causes Enhanced Growth Rate

To determine if the observed changes in GSA activity levels affect the plants morphology and physiology, wild-type and *GSA* over expressing plants were grown in green house conditions and their growth parameters were compared (Fig. 3B). Appreciable differences in general plants growth were observed between transgenic plants and the wild-type control. Total leaf area, total leaf weight, fourth leaf dry weight, Internode third and fourth as well as plant height and specific leaf area were measured and the results showed significance increased in transform plants compared to the wild type (Tabel 1).

**Table 1.** Effect of increased GSA on growth parameter of transgenic lines and the wild type plants.

| **Growth Parameter** | **Total Leaf Area (cm<sup>2</sup>)** | **Total Leaf Weight (gr)** | **Fourth Leaf Dry Weight (gr)** | **Internode Third And Fourth (cm)** | **Plant Height (cm)** | **Specific Leaf Area (SLA)(m<sup>2</sup>.g<sup>-1</sup>)** |
| --- | --- | --- | --- | --- | --- | --- |
| **T0-1** | **591.27±27.33<sup>a</sup>** | **8.98±0.517<sup>a</sup>** | **0.24±0.01<sup>a</sup>** | **0.92±0.076<sup>a</sup>** | **9.5±0.5<sup>a</sup>** | **741.56±30.75<sup>a</sup>** |
| **T0-3** | **573.76±20.84<sup>a</sup>** | **8.9±0.46<sup>a</sup>** | **0.223±0.015<sup>a</sup>** | **0.68±0.076<sup>b</sup>** | **8.4±1.15<sup>a</sup>** | **659.06±20.16<sup>bc</sup>** |
| **T0-4** | **565.35±24.04<sup>a</sup>** | **8.47±0.56<sup>ab</sup>** | **0.22±0.01<sup>b</sup>** | **0.6±0.1<sup>bc</sup>** | **6.73±0.97<sup>b</sup>** | **700.23±25.36<sup>ab</sup>** |
| **Wild Type** | **487.87±17.7<sup>b</sup>** | **7.7±0.3<sup>b</sup>** | **0.2±0.01<sup>b</sup>** | **0.45±0.05<sup>c</sup>** | **6.43±0.51<sup>b</sup>** | **601.63±32.23<sup>c</sup>** |

The values are the mean of five different measurements (n=5). Different letters in the same column represent significant differences at the 95% confidence level.

Increased GSA activity had positive effect on pigment synthesis and the carotenoid contents of transgenic plants.

The data clearly showed that parallel to increase GSA activity in transgenic plants the photosynthesis pigments were significantly altered. The level of Chlorophyll a and b were enhanced significantly from 15 to 110 % of wild type plants (Fig. 4A, 4B). Also, total content of photosynthesis pigment (chlorophyll a and b) was increased 45 to 70 % in transgenic lines (Fig. 4C). The ratio of chlorophyll a and b was reduced significantly 30 to 45% in transgenic plants compared to wild type (Fig. 4D). The amount of carotenoids was also measured from leaves of transgenic and the wild type plants. The results indicated a significant enhanced, 50-65% in carotenoids contents of increased *GSA* lines compared to that from wild type (Fig. 4E).

**Fig. 4.**
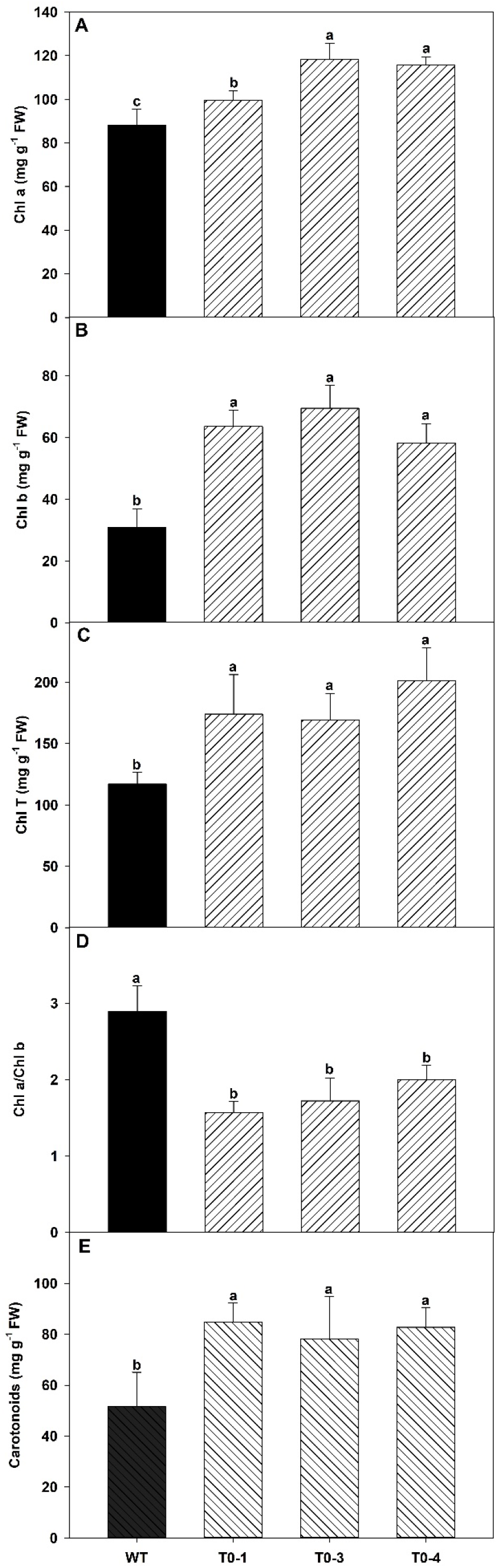
Photosynthesis pigments content of transform lines T0-1, T0-3, T0-4 compared to WT tobacco. (a) chlorophyll total (b) chlorophyll a (c) chlorophyll b (d) Chlorophyll a / Chlorophyll b (e) Carotenoids. Means±SD followed by the same letter(s) in each row are not significantly different, as measured by Duncan’s test (P≤ 0.05). Each data is the average of measurements of 4 plants. Measurements from individual extracts were made in triplicate.

### Over Expression of GSA Activity Elevated Metabolites Include Soluble and Insoluble Sugars as Well as ALA Content in *GSA* Transgenic Lines

To investigate whether increasing GSA activity may correlate with the level of soluble and insoluble sugars, plants materials were sampled from new expanded leaves of *GSA* transgenic lines and wild type plants. The data revealed that the content of soluble sugar was increased in *GSA* over expression lines 60 to 85% about to two fold higher levels than in wild type tobacco plants. The level of insoluble sugars also improved 40 to 115% in T0-1, T0-4 and also in T0-3 respectively compared to that found in the wild type (Fig. 5).

**Fig. 5.**
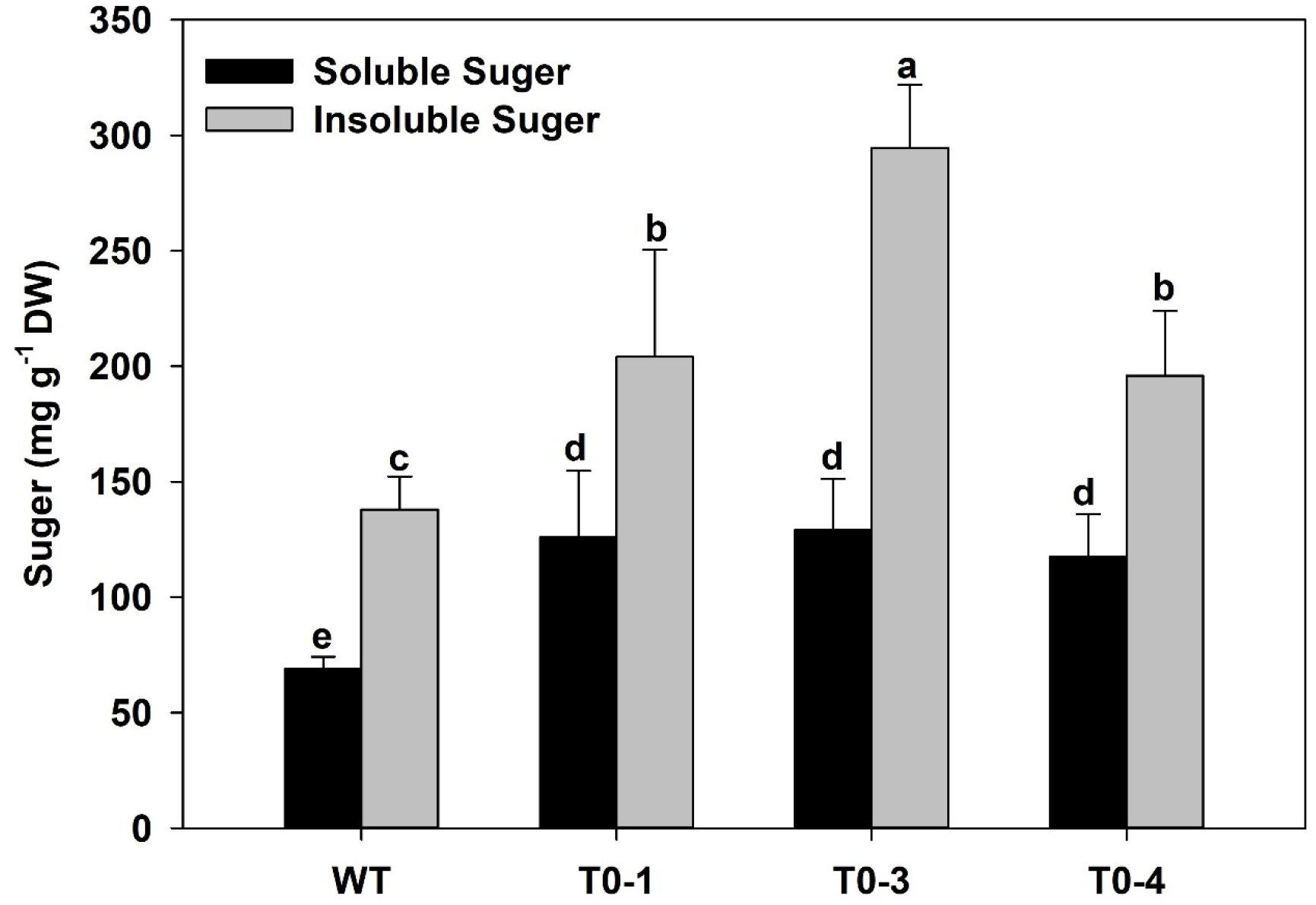
The soluble sugar and insoluble sugar content of fresh leaf of transform lines T0-1, T0-3, T0-4 compared to WT tobacco. Means ± SD followed by the same letter(s) in each row are not significantly different, as measured by Duncan’s test (P≤ 0.05). Each data is the average of measurements of 4 plants. Measurements from individual extracts were made in triplicate.

The data from the previous analysis suggested that chlorophyll and carotenoid contents were increased as *GSA* was expressed in *GSA* over-expressing lines. To test whether increased GSA activity alters the chlorophyll biosynthesis pathway in *GSA* over expression plants, the level of 5-aminolevulinic acid (ALA) was determined. The samples from new expanded leaves of transgenic lines and wild type plant grown in green house conditions were extracted and examined for ALA quantification. As indicated in Fig. 6, the highest significant value for ALA was found in T0-1 (65%), T0-4 (40%) and T0-3 (25%) lines respectively, compared to the equivalent wild type.

**Fig. 6.**
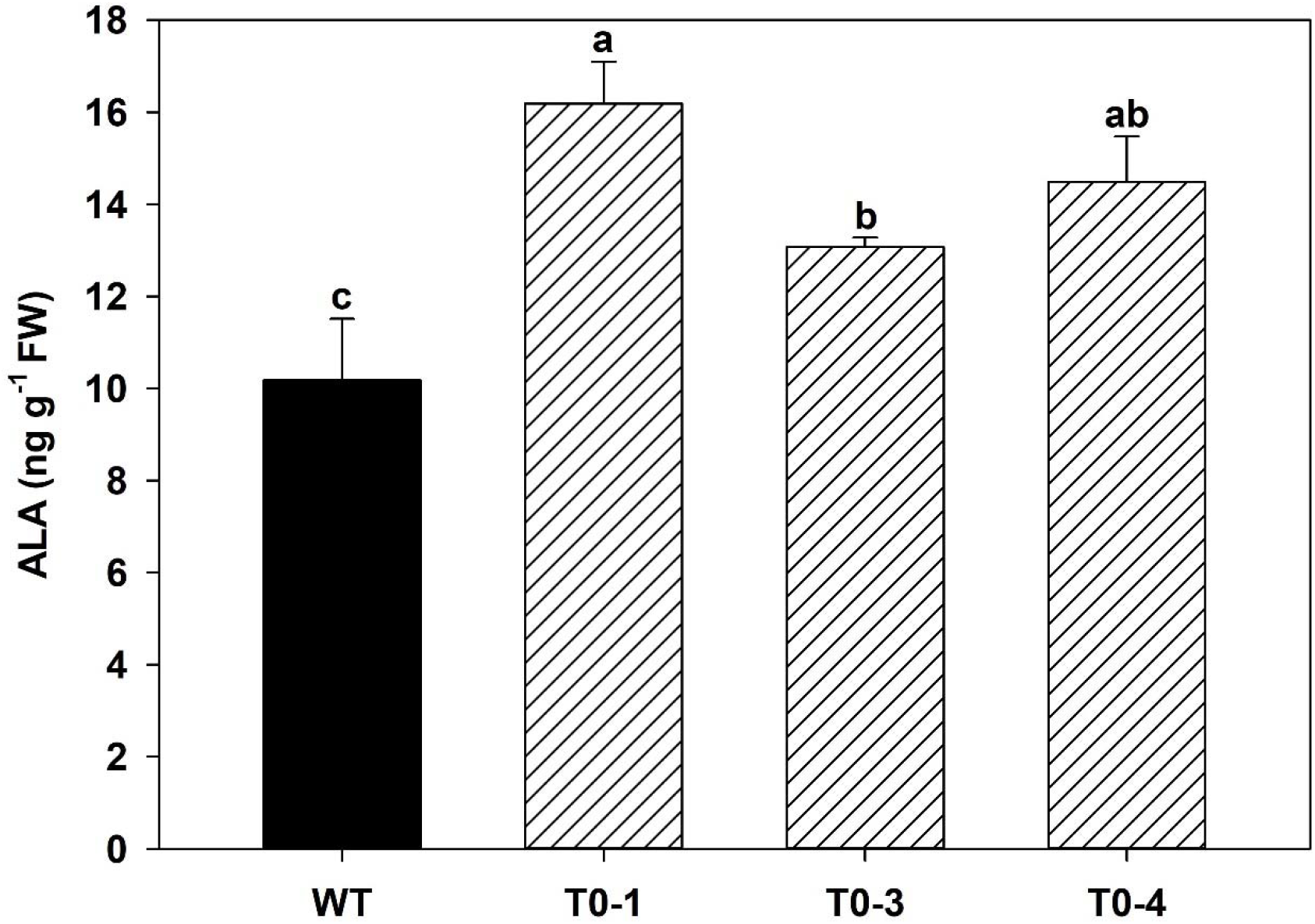
ALA content of fresh leaf of transform lines T0-1, T0-3, T0-4 compared to WT tobacco. Means ± SD followed by the same letter(s) in each row are not significantly different, as measured by Duncan’s test (P≤ 0.05). Each data is the average of measurements of 4 plants. Measurements from individual extracts were made in triplicate.

### Over Expression of GSA Activity Increased Photosynthesis Rate in Transgenic Lines

In order to have a better understanding of increased GSA activity on the function of plant such as photosynthetic rate (A_n_), intercellular CO_2_ concentration (C_i_), plants were grown in greenhouse conditions and the young expanded leaves were subjected to photosynthesis analysis from both transgenic and wild type plants.

Photosynthetic rate was also affected by increasing level of *GSA* in transgenic tobacco plants. As GSA activity increased photosynthetic rate also increased in transgenic plants by 55 to 70% of equivalent wild type plants. High activity of GSA had also positive significant effect on accumulation of internal CO_2_ in transgenic lines (Fig. 7).

**Fig. 7.**
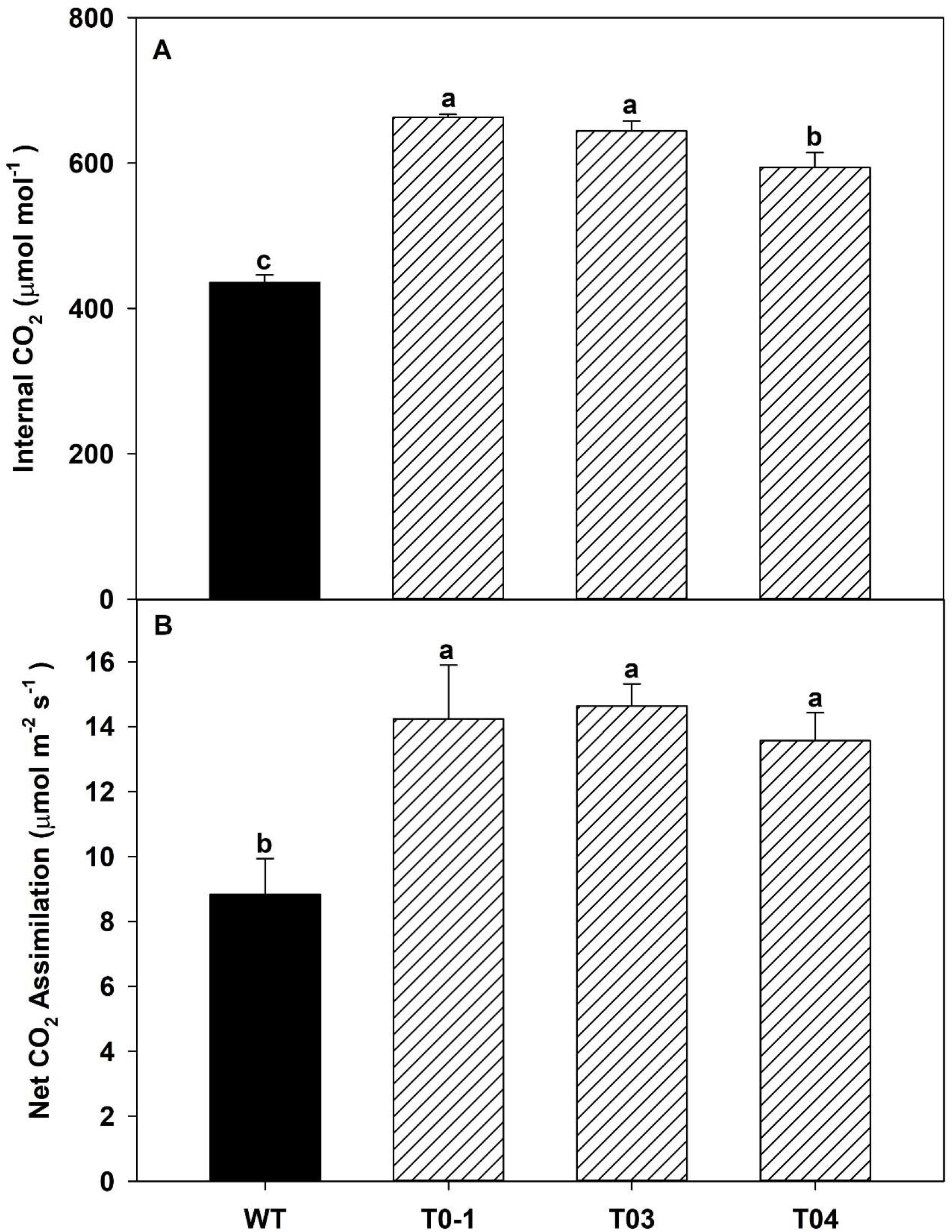
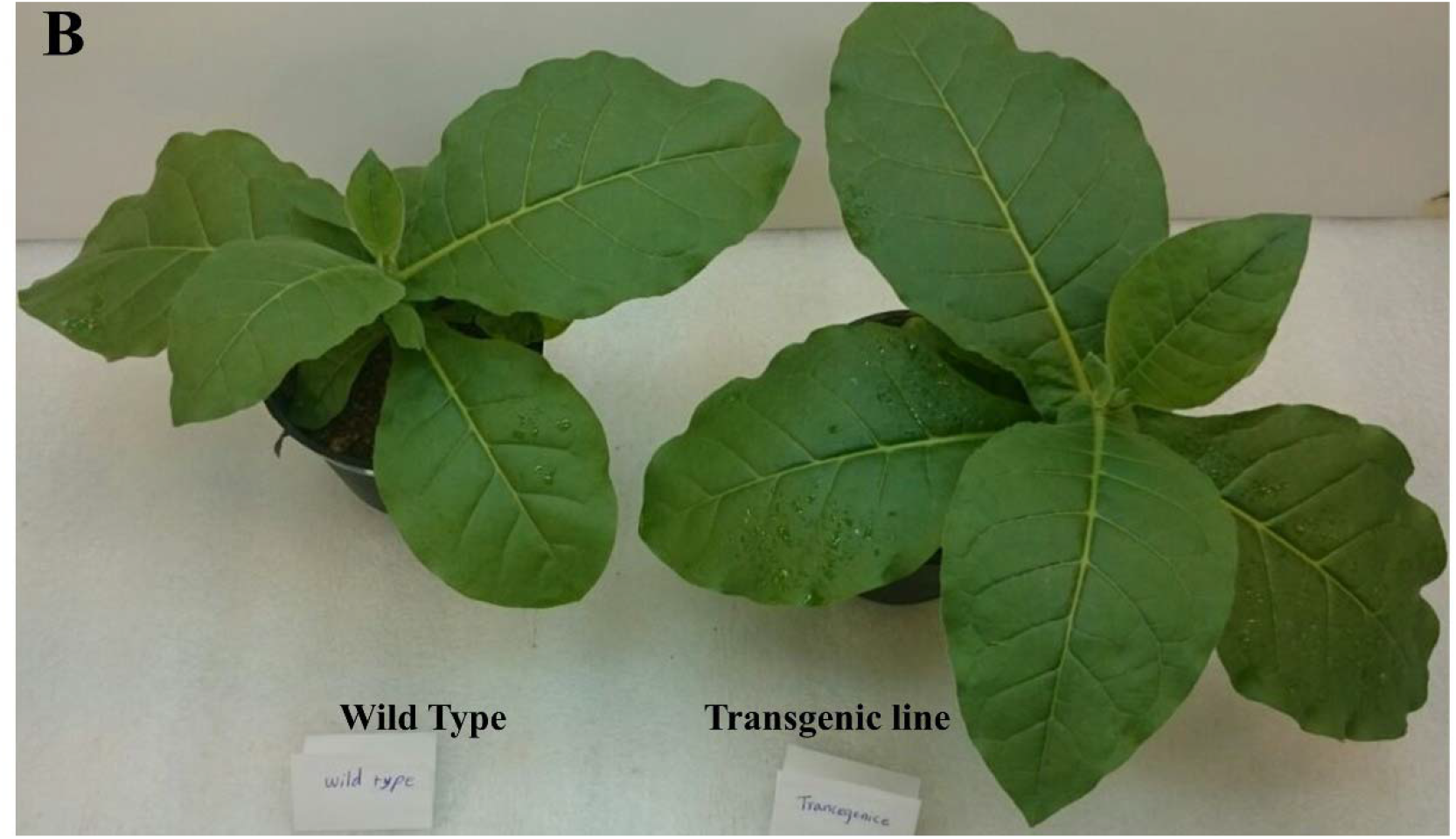
Comparison of leaf photosynthesis parameters between wild type and transgenic. (A) Intercellular CO_2_ concentration; (B) Net CO_2_ Assimilation rate. Means±SD followed by the same letter(s) in each row are not significantly different, as measured by Duncan’s test (P≤ 0.05). Each data is the average of measurements of 4 plants.

## DISCUSSION

We show here that overexpression of *GSA* increases the chlorophyll a and b, total chlorophyll and decreases the Chl a/b ratio in tobacco transgenic lines. Our data also showed that carotenoid content of *GSA* transgenic lines have increased 65% over that of wild type plant. In contrast, Höfgen (1994) claimed that down regulation of GSA activity in transgenic tobacco plants leads to severe plant damage and mimics in some tobacco transformants a wide variety of chlorophyll variegation patterns (Höfgen *et al.*, 1994). Relative to wild type plant, chlorophyll content reduction was varied from 10% to 53% as GSA activity reduced ranging from 22% to 85% in *GSA* antisense plants. Further studies showed that up regulation in activity of chlorophyll biosynthesis enzymes may change the content of chlorophyll and pigments (Höfgen *et al.*, 1994). Biswal et al. (2012) reported that over expression of *chlorophyllide a oxygenase* resulted in a high level of chlorophyll b but, Chl a/b ratio was decreased in tobacco or Arabidopsis *CAOx* plants were grown in both low or high light conditions (Tanaka *et al.*, 2001; Biswal *et al.*, 2012). Pattanayak et al. (2011) also reported that over expression *protochlorophyllide oxidoreductase C (PORC)* had increased Chlorophyll content up to 28% over that of WT plants and had also a little higher carotenoid content (Pattanayak and Tripathy, 2011). In our study, simultaneous increased of ALA synthesis rate was observed in response to the elevation in activity of GSA enzyme in *GSA* over expression tobacco plants. It is worth mentioning that increased in capacity of ALA synthesis was considered as a main factor to improve the pigment and chlorophyll content in transgenic lines. This was highly due to the function of ALA as a precursor to form the synthesis of chlorophylls, heme, and phycobilins in plants, algae, and many bacteria (Kannangara and Gough 1978; Sangwan and O’Brian 1993). The data obtained from ALA supplementation also suggests that exogenous ALA increased the content of chlorophylls, tetrapyrols and consequently promotes photosynthesis rate (Tanaka *et al.*, 1993; Hotta *et al.*, 1997; Xu *et al.*, 2011; Ye *et al.*, 2016; Tang *et al.*, 2017; Liu *et al.*, 2018).

In *GSA* over expression plants grown in greenhouse conditions, starch and soluble sugars content substantially increased to two fold as compared with that in wild-type plants throughout the diurnal cycle. Preliminarily analysis of *GSA* over expression lines using iodine staining also revealed (data not shown) that starch remained at the end of the day while no starch was detectable at the beginning of the day in both transgenic and wild type plants. These data provide evidence that synthesis or degradation of starch in transgenic plants over the day/night cycle was completely normal and accumulation of starch was most likely affected by increased activity of GSA rather than to extend any other reason in *GSA* over expression lines. These results showed that the additional starch produced in the *GSA* transgenic plants was being used in the dark period, possibly for plant growth and this was observed in increased of leaf area, dry weight and plant height.

Our data, along with those reported by Lefebvre et al. (2005), revealed a clear correlation between increased photosynthesis and starch accumulation (Lefebvre *et al.*, 2005). Indeed, the total leaf weight of greenhouse-grown *GSA* plants increased by up to 10 to 17%, as compared with wild-type plants. This finding is in agreement with data from a previous report in plants overexpressing. Sedoheptulose-bisphosphatase enzyme, where PSII photosynthetic efficiency, carbon assimilation, starch level, and dry matter accumulation consistently increased (Lefebvre *et al.*, 2005). Along the same line, Transgenic tobacco plants over-expressing a chloroplast-targeted FBPase/SBPase bifunctional cyanobacterial enzyme also showed a 24% increase in final dry matter and a 50% increase in photosynthetic CO_2_ fixation (Miyagawa *et al.*, 2001). It can be postulated that some of the enzyme involved in the Calvin cycle are in excess amount and reduction up to 40% in activity of this enzymes compared to wild type have no dramatic effect on photosynthesis capacity whereas, photosynthesis is very sensitive with a small reduction on activity of Transketolase and SBPase (Quick *et al.*, 2004; Rosenthal *et al.*, 2011; Khozeai *et al.*, 2015). Small reduction in plastid transketolase activity (20% to 40%) reduced the supply of ribulose-1, 5-bisphosphate (important substrate to regenerate third phase of Calvin cycle) and resulted in an inhibition of photosynthesis (Henkes *et al.*, 2001). Furthermore, Chlorophyll was unaltered in Transketolase antisense plants with 50% less in activity of transketolase compared to wild type. However, greater inhibition of TK expression led to localized loss of chlorophyll along the leaf. Total carotenoids also decreased when TK activity decreased to 50% of the wild-type value (Henkes *et al.*, 2001). In contrast, over expression of transketolase in TK overexpressing tobacco plants changed the carbon flux with negative effect on thiamin biosynthesis pathway and extending a chlorotic phenotype accompanied with retardation in growth due to the shortage of cofactor thiamin pyrophosyhate for the TPP dependent enzymes (Khozeai *et al.*, 2015). The same characters as photosynthesis, chlorophyll, carotenoids and growth rate were reported in antisense SBPase tobacco plant, reductions in SBPase activity resulted in a decline in the rate of carbon assimilation in the antisense plants, and this reduction was higher under saturating CO_2_ (Harrison *et al.*, 1998). The chlorophyll content of the leave of antisense SBPase plants was similar in wild-type except those plants with reduction of SBPase activity less than 20% of wild-type. This response resembles that seen in transformants with decrease expression of *glutamate-1-semialdehyde aminotransferase* (Höfgen *et al.*, 1994).

This was also confirmed that decreased TK and SBPase expression resulted a reduction in amount of shoot fresh weight, dry weight, leaf area and a marked decrease in shoot length and shoot biomass. The diurnal turnover of sucrose and starch in TK antisense showed a significant decreased in the content of carbohydrate levels such as sucrose, glucose and fructose in the leaves, whereas starch remained at high level (Harrison *et al.*, 1998).

Another interesting observation in our study was related to photosynthesis rate in *GSA* over expression lines. There was an additional increase (40-50%) in net assimilation rate of *GSA* over expression lines compared to wild type plants. Our results are similar to those reported by Biswal et al. (2012) when the activity of chlorophyllide a oxygenas increased in CAO-overexpression tobacco and Arabidopsis plants and elevation of ALA, accumulation of chlorophyll content and increased in photosynthesis rate were evident. The previous finding data provide evidence that increased in photosynthesis rate in chlorophyllide a oxygenas overexpressed plants was mainly due to increase in chlorophyll b synthesis which was accompanied by an increased in light-harvesting chlorophyll proteins and light absorption level (Tanaka *et al.*, 2001; Biswal *et al.*, 2012). Our finding confirmed that any change in activity of enzymes in chlorophyll biosynthesis pathway caused the variation in content of chlorophylls and consequently altered the photosynthesis rate (Papenbrock *et al.*, 2000; Tanaka *et al.*, 2001; Shalygo *et al.*, 2009). We hypothesis that photosynthesis is very sensitive to a small reduction/expression on activity of enzymes in chlorophyll biosynthesis pathway and this was mainly due to the mechanism of light absorbing and energy production occurred in this pathway. Altogether made the chlorophyll biosynthesis pathway as a great target for genetic modification leading to increase photosynthesis and for future improvement in crop productivity.

## ACKNOWLEDGMENTS

We would like to express our thankful to RNA biotechnology Company for their vast input throughout this project.

## Abbreviations

ALA: 5-aminolevulinic acid
GSA: *Glutamate-semialdehyde aminotransferase*

